# EcDBS1R6: A new broad-spectrum cationic antibacterial peptide derived from a signal peptide sequence

**DOI:** 10.1101/869867

**Authors:** William F. Porto, Luz N. Irazazabal, Vincent Humblot, Evan F. Haney, Suzana M. Ribeiro, Robert E. W. Hancock, Ali Ladram, Octavio L. Franco

## Abstract

Bacterial infections represent a major worldwide health problem, with an special highlight on Gram-negative bacteria, which were assigned by the World Health Organization (WHO) as the most critical priority for development of novel antimicrobial compounds. Antimicrobial peptides (AMPs) have been considered as potential alternative agents for treating these infections. Here we demonstrated the broad-spectrum activity of EcDBS1R6, a peptide derived from a signal peptide sequence of *Escherichia coli* that we previously turned into an AMP by making changes predicted through the Joker algorithm. Signal peptides are known to naturally interact with membranes; however, the modifications introduced by Joker made this peptide capable of killing bacteria. Membrane damage of the bacterial cells was observed by measuring membrane integrity using fluorescent probes and through scanning electron microscopy imaging. Structural analysis revealed that the C-terminus was unable to fold into an α-helix, indicating that the EcDBS1R6 antibacterial activity core was located at the N-terminus, corresponding to the signal peptide portion of the parent peptide. Therefore, the strategy of transforming signal peptides into AMPs seems to be promising and could be used for producing novel antimicrobial agents.

## 1 Introduction

The discovery and development of antimicrobial agents made a number of infectious diseases treatable and curable. However, today, 90 years after the discovery of penicillin by Sir Alexander Fleming, the threat of bacterial infections has become major (1). This is a consequence of two factors, namely bacterial resistance development to conventional antimicrobial agents and the small number of new antimicrobial compounds, and especially new structural classes, that have reached the market in the past decade (1). Recently, 30 million sepsis cases, for which the frontline treatment is antibiotics, are estimated to occur worldwide each year, resulting in potentially 5-8 million deaths (2). In this alarming situation, the World Health Organization (WHO) released a list of bacterial pathogens that require the development of new antimicrobial agents, including *Acinetobacter baumannii*, *Pseudomonas aeruginosa* and several Enterobacteriaceae in their critical priority list (3).

Among potential new drugs, the antimicrobial peptides (AMPs) have gained attention, especially those having broad-spectrum activity and no or low toxicity for mammalian cells (4–6). In general, they have an amphipathic and cationic structure, with a net positive charge between +3 and +9, and their size can range from 12 to 50 amino acid residues (6). Despite their overall hydrophobic and cationic character, the combinatorial space for peptide design is enormous (i.e. for a 20 mer peptide, there are 20^20^ possibilities). Therefore a number of computational methods have been applied to design novel AMP sequences (7–10), as the initial step in the development of new antimicrobial compounds, followed by in-depth characterization of the lead molecules.

Recently, our group developed the Joker algorithm for designing novel AMPs (8). This algorithm works in a sliding window fashion, using specific substitution rules to imprint an antimicrobial pattern into a target sequence, with an overall success rate of 64% (8). Joker provides an algorithm that is an initial step in developing novel AMPs and 84 sequences were screened in our initial study. Now we have started the second step of characterizing the lead peptides (11, 12).

Here we selected the peptide EcDBS1R6 (Figure 1) for an in-depth characterization of its antimicrobial spectrum and shed some light regarding its mechanism of action. EcDBS1R6 was designed using Joker applied to a signal peptide sequence from *Escherichia coli* (Figure 1) (8). Signal peptides naturally interact with cell membranes, having the capacity to insert spontaneously into phospholipid bilayers (13–15). Thus, with the modifications introduced by Joker, the sequence was shown to be able to kill bacteria by targeting their membranes. Together with an analysis of cytotoxicity, we were able to demonstrate that the peptide has a reasonable selectivity for microorganisms of the WHO “priority pathogens” list. Analysis of the EcDBS1R6-killing kinetics and bacterial mechanism revealed that the peptide is able to rapidly damage the bacterial membrane. The effects on bacterial morphology were visualized by FEG-SEM imaging. Moreover, the peptide’s three-dimensional structure was evaluated by using 50 independent runs of molecular dynamics simulations.

**Figure 1.**
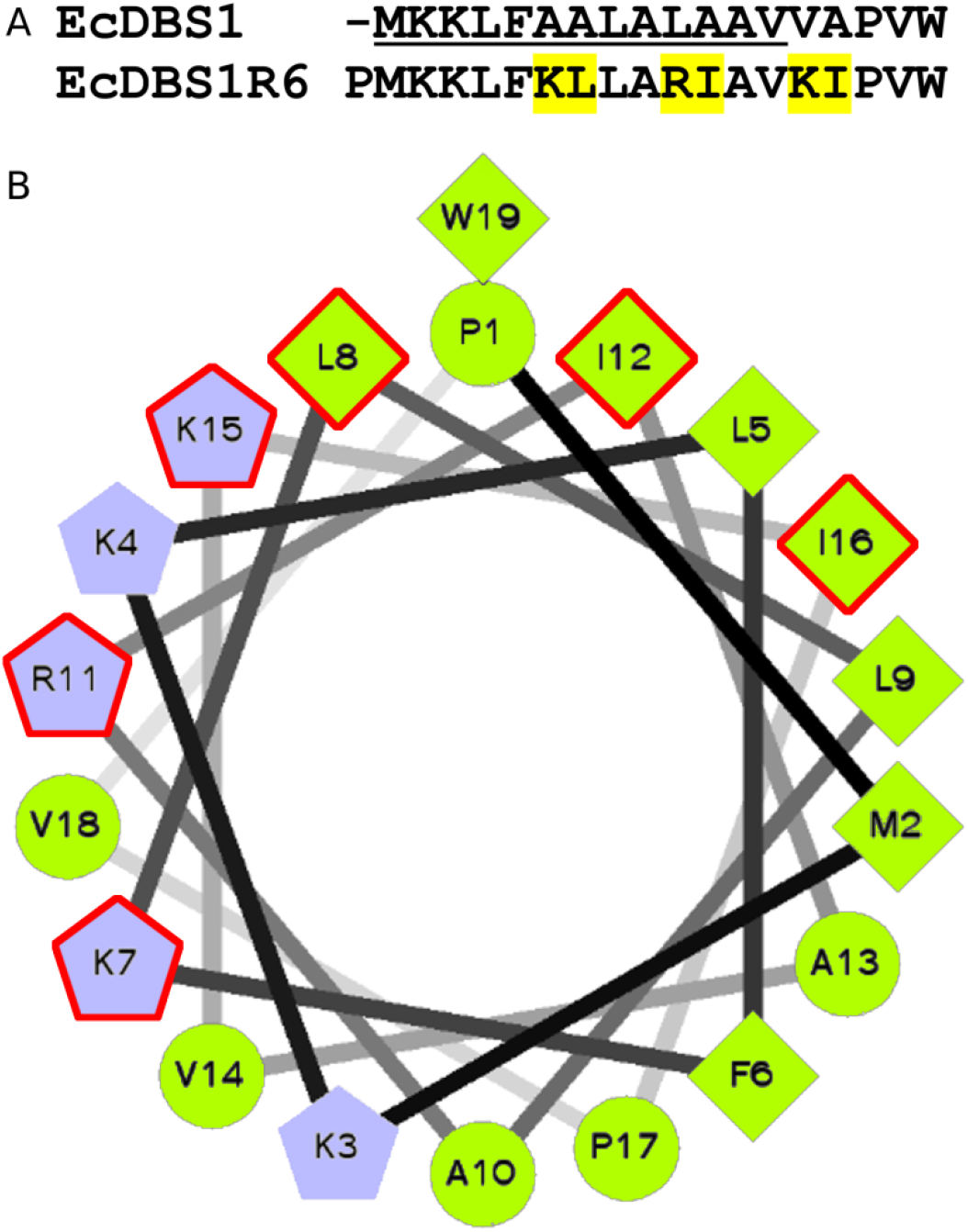
Design and prediction of EcDBS1R6. (A) Sequence alignment of EcDBS1R6 and its parental sequence corresponding to the N-terminus fragment of a mercury transporter protein from *Escherichia coli* (GenBank ID CAA70241). In addition to the N-terminus capping with a proline residue, EcDBS1R6 was generated by modifying the sixth sliding window using the antimicrobial pattern “KL[AL]x[RKD][ILV]xxKI”. Amino acids between brackets represent positions that could be filled up by one of the indicated residues whereas “x” represent positions that could be filled up by any of the 20 natural amino acid residues. The phobius prediction (13) indicated a signal peptide topology for the original sequence is underlined. (B) Schiffer-Edmundson helical wheel projection of EcDBS1R6 (http://rzlab.ucr.edu/scripts/wheel/wheel.cgi). Hydrophobic residues are represented by green circles (tiny side chains) or green diamonds (large side chains), while the positively charged residues in blue pentagons; the mutated residues are outlined in red. The designed sequence displayed 63% hydrophobicity and a net charge of +5, being predicted to assume an amphipathic α-helix conformation. EcDBS1R6 showed a MIC of 6.25 μg.mL^−1^ in the screening against a bioluminescent strain of *Pseudomonas aeruginosa*, and exhibited no hemolytic activity against human erythrocytes (8).

## 2 Results

### 2.1 EcDBS1R6 displays broad-spectrum activity

Minimal inhibitory concentrations (MICs) of EcDBS1R6 were determined against several bacterial strains and yeasts. As indicated in Table 1, MIC values demonstrated that EcDBS1R6 displayed potent broad-spectrum antibacterial activity at low concentrations, in the range of 3 – 6.25 μM, against Gram-negative bacteria. Against Gram-positive bacteria, EcDBS1R6 was also highly active with MICs in the range of 3 – 25 μM, except *E. faecalis* (MIC ≥ 100 μM). EcDBS1R6 was found to be inactive against *Candida* yeast species, such as *C. albicans* and *C. parapsilosis* (MIC = 100 μM, Table 1).

**Table 1.** Antimicrobial activity and cytotoxicity of EcDBS1R6.

| Cell strain | Microorganism/cell line | Active concentration (μM)* |
| --- | --- | --- |
| **Gram-negative bacteria** | *Escherichia coli* ATCC 25922 | 3 |
|  | *Pseudomonas aeruginosa* ATCC 27853 | 6.25 |
|  | *Klebsiella pneumoniae* ATCC 13883 | 3 |
|  | *Acinetobacter baumannii* ATCC 19606 | 3 |
| **Gram-positive bacteria** | *Staphylococcus aureus* ATCC 25923 | 3 |
|  | *Streptococcus pyogenes* ATCC 19615 | 25 |
|  | *Listeria ivanovii* Li 4pVS2 | 12.5 |
|  | *Enterococcus faecalis* ATCC 29212 | > 100 |
| **Yeasts** | *Candida albicans* ATCC 90028 | 100 |
|  | *Candida parapsilosis* ATCC 22019 | 100 |
| **Human cells** | HEK-293 | 111.1 ± 27.2 |

\*Minimum inhibitory concentrations (MICs) for microorganisms and inhibitory concentration 50 (IC<sub>50</sub>) for HEK-293 human cells are expressed as average values from three independent experiments performed in triplicate.

### 2.2 EcDBS1R6 has a safe in vitro selectivity index

We examined the cytotoxicity of EcDBS1R6 toward human embryonic kidney-293 cells (HEK-293), since the kidney plays an important role in the systemic clearance of administered drugs. The results contained in Table 1 indicated that EcDBS1R6 is not cytotoxic against HEK-293 cells at the antimicrobial concentrations (IC_50_ = 111.1 ± 27.2 μM). Taking into account the MICs against Gram-negative bacteria and the cytotoxicity assessments, EcDBS1R6 showed a selectivity index of 12.92, indicating that to achieve a toxic effect a 13-fold higher administration of this peptide would be necessary.

### 2.3 EcDBS1R6 acts through a membranolytic mechanism

The killing effect of EcDBS1R6 was investigated on Gram-negative and Gram-positive bacteria using the reference strains *E. coli* (ATCC 25922) and *S. aureus* (ATCC 25923), respectively, at concentrations 2-fold above the MIC (6.25 μM). The time-curves revealed that EcDBS1R6 caused a rapid complete killing of both *E. coli* and *S. aureus* cells within the first 15 minutes (Figure 2). The potent effect of EcDBS1R6 observed against these reference strains (MIC = 3 μM, Table 1) was therefore bactericidal.

**Figure 2.**
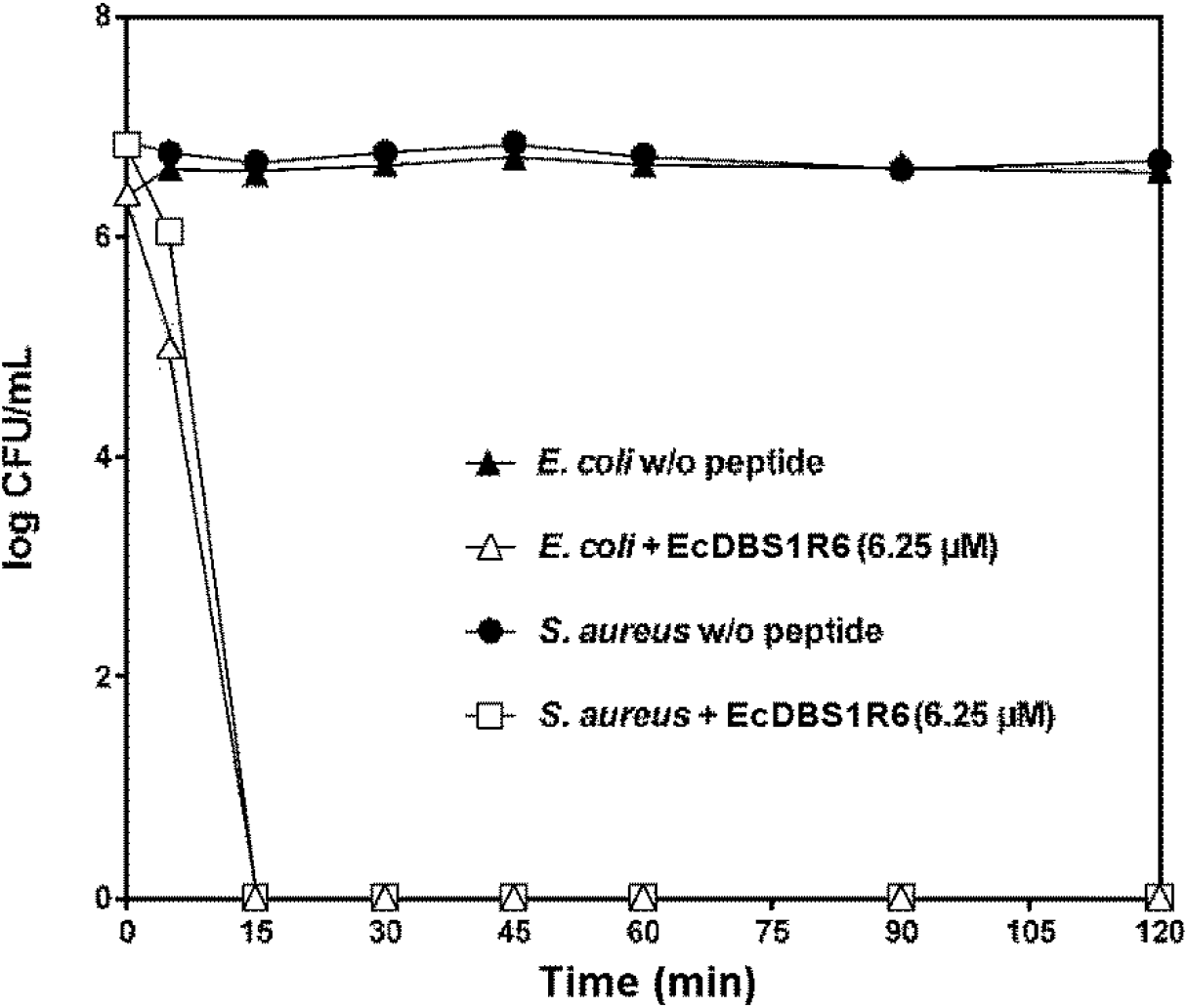
Time-dependent killing of *E. coli* ATCC 25922 and *S. aureus* ATCC 25923 induced by 6.25 μM EcDBS1R6 (2 × MIC). Controls correspond to bacteria incubated in PBS without peptide. The values are the means ± S.E.M. of one representative experiment performed in triplicate.

To gain insight into the effect of EcDBS1R6 on the bacterial cytoplasmic membrane, we used the two fluorescent dyes, SYTOX Green (SG) and DiSC3(5), to investigate the ability to induce membrane permeabilization and loss of membrane potential, respectively. The two reference strains, *E. coli* ATCC 25922 and *S. aureus* ATCC 25923, were treated with a concentration 2-fold above the MIC (6,25 μM), using the membrane permeabilizing bee venom peptide melittin (5 μM) as a positive control. In Figure 3, the specific increase of the SG fluorescence (compared to the negative control) observed after incubation of the bacteria with EcDBS1R6 revealed that the peptide was able to permeabilize the cytoplasmic membrane of both *E. coli* (Figure 3A) and *S. aureus* (Figure 3B). For *E. coli*, after a short lag time, the complete permeabilization induced by EcDBS1R6 was as efficient as melittin. For *S. aureus*, a rapid permeabilization occurred with a maximum SG fluorescence signal lower than the positive control. Regarding the depolarization of membrane potential, an immediate effect on *E. coli* (Figure 3C) and *S. aureus* (Figure 3D) was observed, as indicated by the increase in the DiSC3(5) fluorescence by the peptides when compared to the PBS negative control) For both bacterial cells, in the presence of EcDBS1R6, a rapid and potent depolarization was achieved with a maximal threshold equal (*E. coli*) or similar (*S. aureus*) to the positive control, melittin.

**Figure 3.**
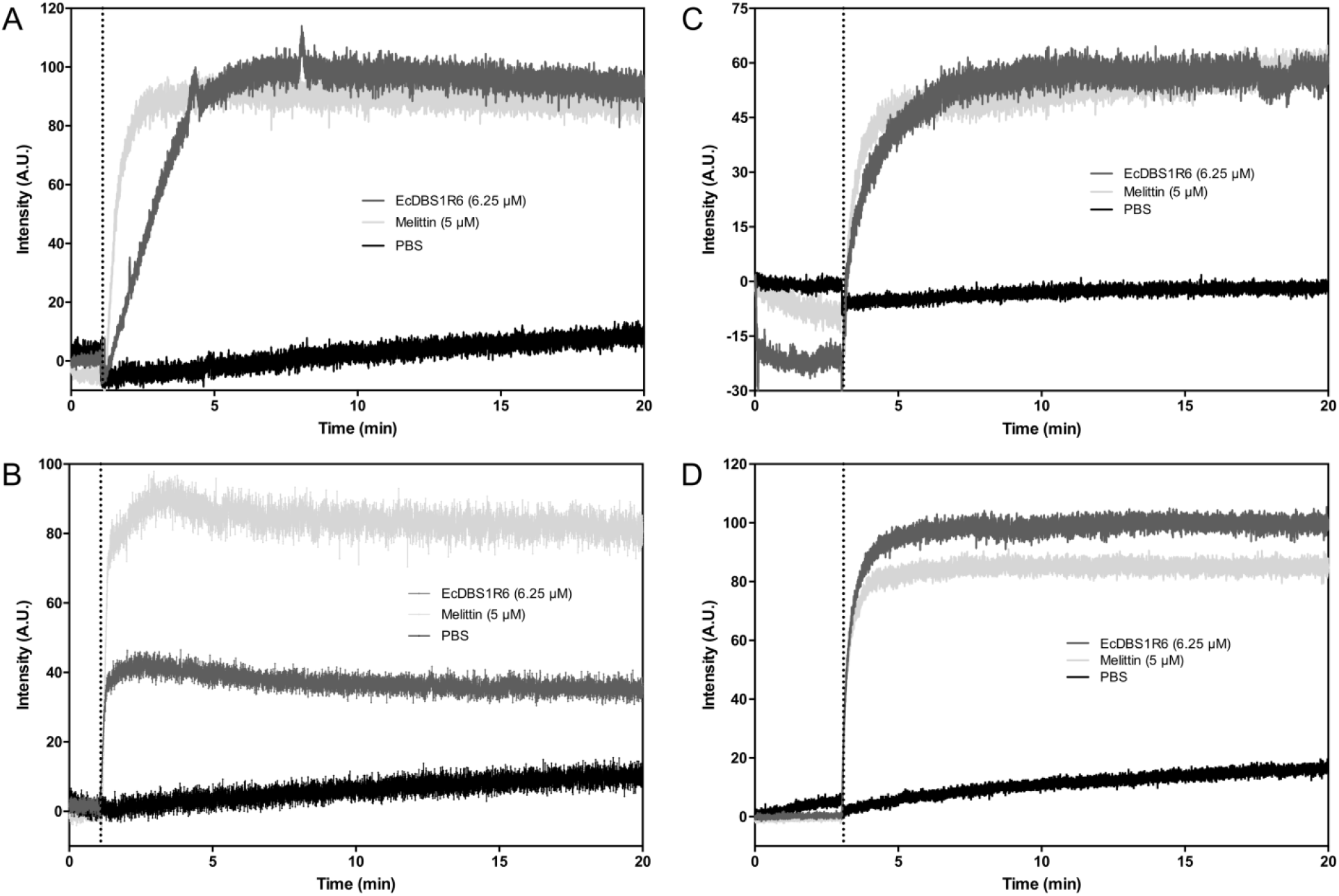
Bacterial membrane permeabilization (A, B) and depolarization (C, D) after incubation with EcDBS1R6. The kinetics of membrane permeabilization of *E. coli* **(A)** and *S. aureus* **(B)** was determined by SYTOX green uptake assay, after addition. **(C-D)** Effect of EcDBS1R6 on the cytoplasmic membrane potential of *E. coli* **(C)** and *S. aureus* **(D)** followed by monitoring the fluorescence of DiSC3(5) dye. The negative control (PBS) corresponds to the bacteria incubated without peptide, whereas the positive control corresponds to the incubation with melittin 5 μM. The vertical dotted line indicates peptide (or PBS) addition.

To visualize the antibacterial activity of EcDBS1R6, FEG-SEM investigations were performed on the Gram-negative bacterium *P. aeruginosa* and the Gram-positive bacterium *L. ivanovii* at peptide concentrations 2-fold above the lethal concentration. Figure 4 clearly shows that all the bacterial cells suffered alterations in membrane integrity after exposure to EcDBS1R6. The high activity of EcDBS1R6 against *P. aeruginosa* was confirmed by the observation of membrane deformation after contact with 12.5 μM of peptide (Figure 4B), as compared to untreated cells (Figure 4a). For *L. ivanovii*, Figure 4 revealed also membrane damage after treatment with 25 μM EcDBS1R6 (Figure 4D) when compared to intact cells (Figure 4C).

**Figure 4.**
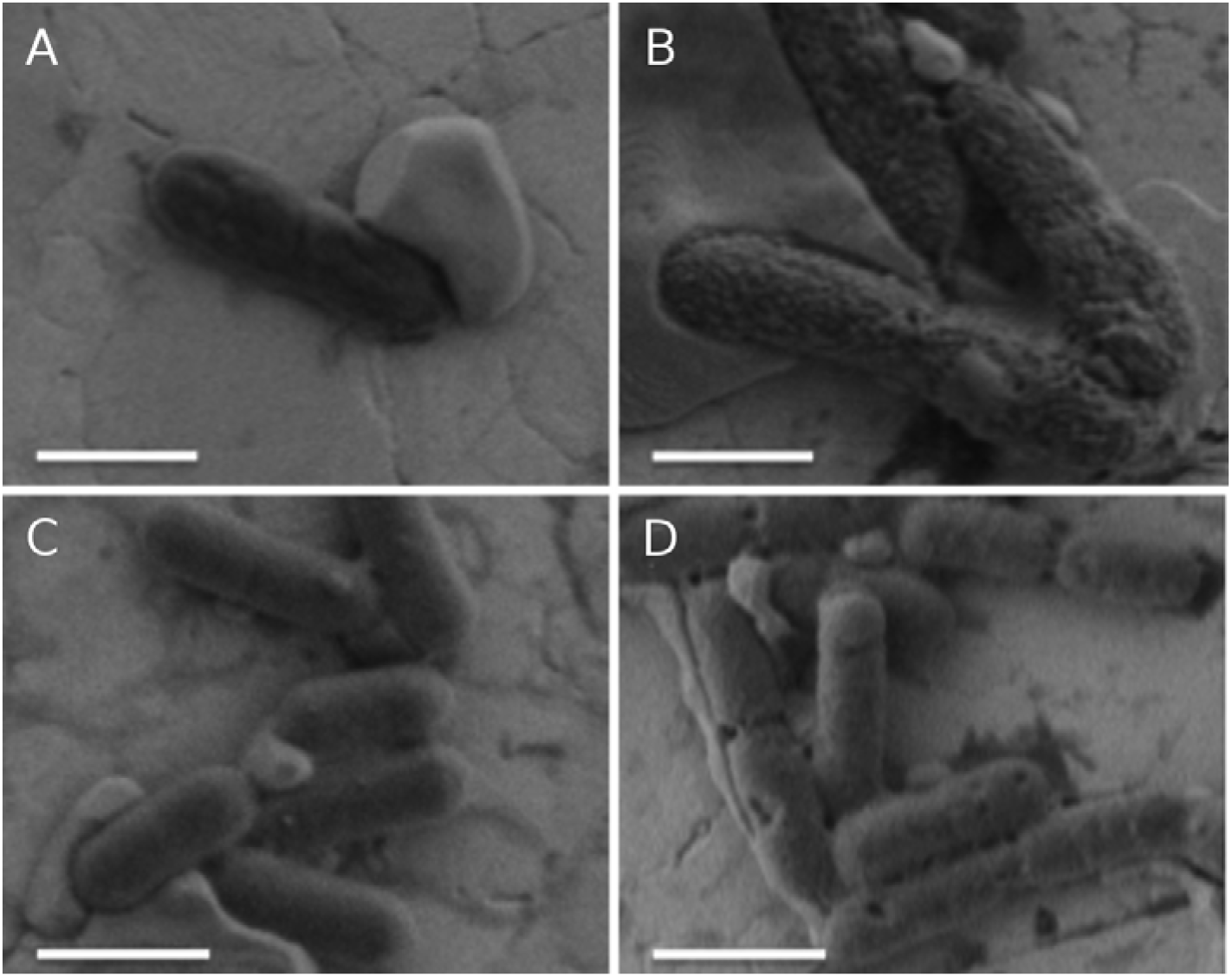
FEG-SEM visualization of EcDBS1R6 effects on Gram-negative and Gram-positive bacterial cells. (A) *P. aeruginosa* ATCC 27853 without peptide. (B) *P. aeruginosa* treated with 12.5 μM EcDBS1R6 (concentratio 2-fold above the MIC). (C) *L. ivanovii* Li 4pVS2 without peptide. (D) *L. ivanovii* treated with 25 μM EcDBS1R6 (concentration 2-fold above the MIC). Scale bar = 1μM.

### 2.4 The N-terminus is the core of EcDBS1R6 activity

To determine the core of EcDBS1R6 activity, we applied a strategy commonly used for high-throughput screening to scan the peptide sequence. Ten fragments synthesized on a cellulose membrane support, derived based on a sliding window of ten residues, were assessed against a bioluminescent strain of *P. aeruginosa*. The results revealed that the first four N-terminus fragments were the most active, with MICs of 12.5 or 25 μg.mL^−1^ (Figure 5). Also, the three C-terminus residues seemed to not contribute to the activity, since the fragments harboring them were inactive (MIC>200 μg.mL^−1^) (Figure 5).

**Figure 5.**
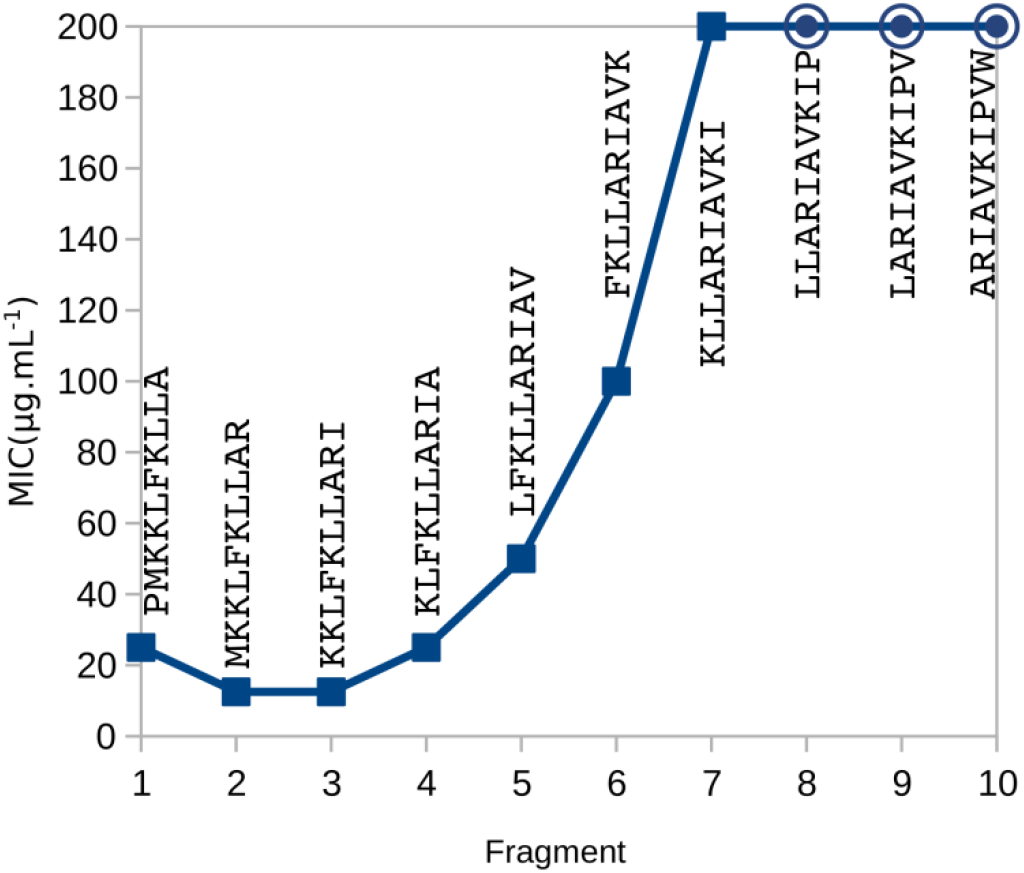
EcDBS1R6 sliding window scanning. The sequence was subjected to a sliding window system of te residues, generating ten fragments, which were synthesized in cellulose support by means of the SPOT technology and then, were evaluated against a bioluminescent strain of *P. aeruginosa*. The first four fragments displayed the lower MICs (25 or 12.5 μg.mL^−1^), while the last three were inactive at the highest concentration tested (200 μg.mL^−1^) and thus, they are marked with circles. The sequence for each fragment is displayed near to its respective marker.

### 2.5 The C-terminus is pivotal for EcDBS1R6 structure

As previously indicated (8), the three-dimensional structure of EcDBS1R6 was predicted in vacuum to be composed of an α-helix comprising the residues Met^2^-Ile^16^, while the residues Pro^17^-Trp^19^ were proposed to present an extended conformation that was folded back over the hydrophobic counterpart (data not shown). To analyze the structure in a membrane-mimetic environment, a series of 50 molecular dynamics simulations were performed in 50% of TFE (v/v). Simulations indicated that there are four main conformations (Figure 6). The extended conformation, where the peptide underwent a complete unfolding process was rare, while the maintenance of the α-helix was more frequent (Figure 6).

**Figure 6.**
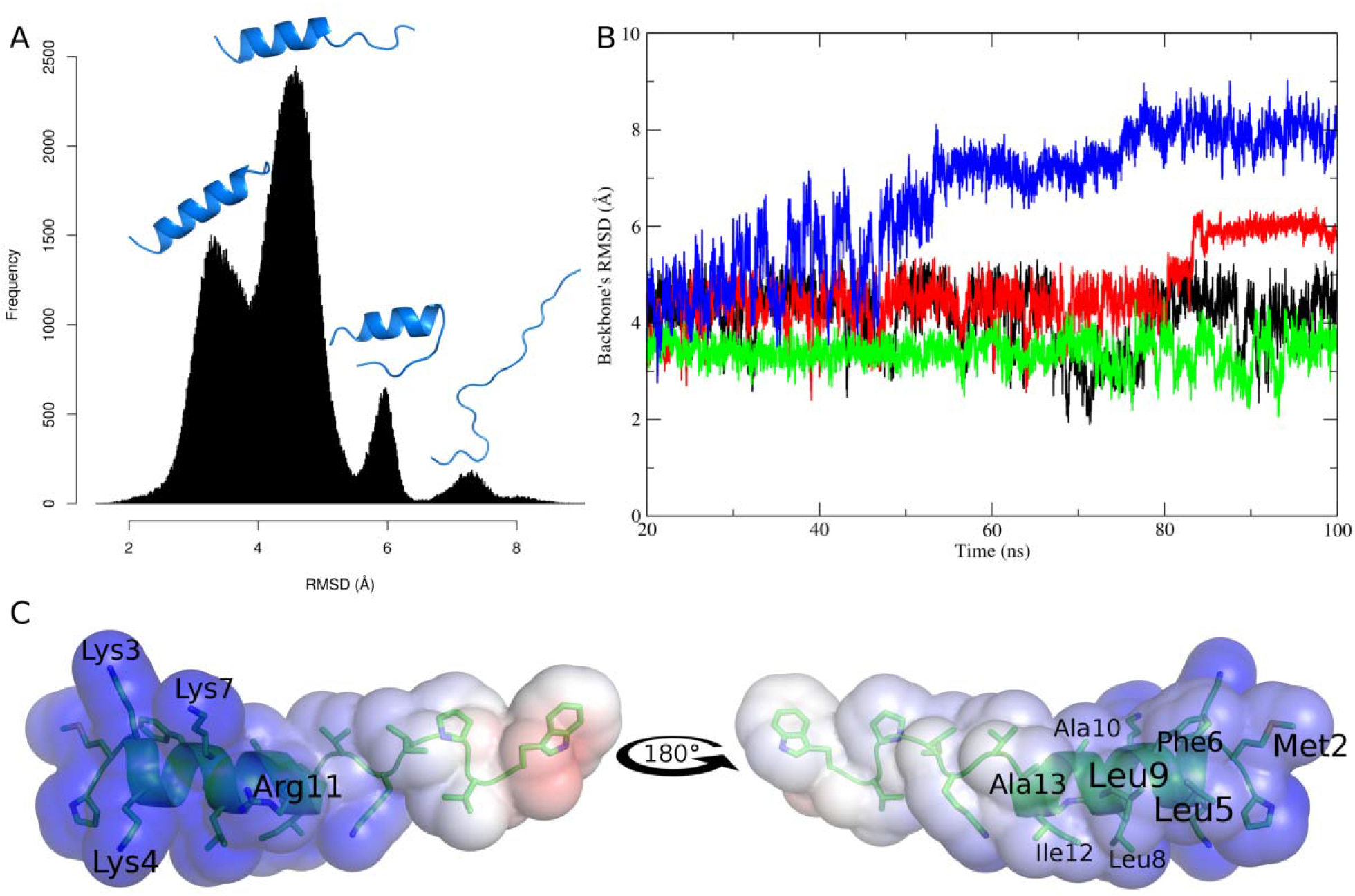
*In silico* structural analysis of EcDBS1R6. The *ab initio* model of EcDBS1R6 was subjected t 50 independent runs of molecular dynamics simulations in a water-TFE mixture (50%, v/v). (A) RMSD histogram considering the interval between 20 and 100 ns. In such period, a quadrimodal distribution could be observed. A representative structure is displayed for each mode. (B) Four representativ trajectories for each structural type, the blue trajectory indicated the unfolded structure; the red one, th structure with the C-terminus folded back to the α-helix; the black one corresponds to the structure with extended C-terminus loop; and the green one, the complete α-helix structure. (C) EcDBS1R6 electrostatic surface. Surface potentials were set to ±5 kTe^−1^ (133.56□mV). Blue indicates positively charged regions, red the negatively charged regions and white non-polar ones.

The C-terminus played a pivotal role in the peptide conformation. The initial C-terminal loop was spread from Pro^17^-Trp^19^ to Ala^12^-Trp^19^ residues, and appeared as an extended conformation (which is the most common) or folded back over the α-helix (Figure 6). Also there was a possibility to maintain the initial conformation (Figure 6), which is the second most common conformation. However, due to the presence of Pro^17^, which is known to be a α-helix breaker, this initial α-helical conformation was not stable. Based on the most frequent conformation, the electrostatic surface was calculated. The extended region was essentially hydrophobic, even with the presence of Lys^15^ (Figure 6C), while the α-helix portion presented a well-defined amphipathic structure with a cationic cluster composed of Lys^3^, Lys^4^, Lys^7^ and Arg^11^ (Figure 6C), and a hydrophobic cluster composed of Met^2^, Leu^5^, Phe^6^, Leu^8^, Leu^9^, Ala^10^, Ile^12^ and Ala^13^ (Figure 6C).

## 3 Discussion

The need for novel antimicrobial agents is undeniable and has caused AMPs to be more extensively evaluated as drugs of the future. In fact, the application of computational methods for AMPs in the last years has enabled the field to move forward, both in identification of natural peptides (5) and through computer-aided design of synthetic peptides (4, 16). The application of such algorithms allows exploration of new amino acid combinations, which in turn, could lead to solutions to the need for new molecules to threat bacterial infections. The peptide EcDBS1R6 was developed using the Joker algorithm (8), a recent method that extrapolates the linguistic model for rational design of antimicrobial peptides (17) to a card game model, combining peptides sequences with antimicrobial patterns.

The first prototypes indicated that the peptides designed using Joker were selective against Gram-negative bacteria (8), which are amongst those with the highest priority on the WHO list (3). However, as new peptides were tested, an ability to kill Gram-positive bacteria could be observed, e.g. for PaDBS1R1 (11) and EcDBS1R5 (12). EcDBS1R6 (18) is also capable of killing Gram-positive bacteria, in addition to Gram-negatives (Table 1). Together with the absence of toxicity at antimicrobial concentrations, this peptide had a selective index of 12.92 against Gram-negative bacteria, which allows it to be considered as a safe drug, according to the U.S. Food and Drug Administration (19), especially due to its absence of toxicity to kidney cells (Table 1).

Despite the absence of peptides with similar sequences in AMP databases (8), the EcDBS1R6 sequence is composed of lysine residues, as the cationic portion, and aliphatic residues, including leucine and isoleucine, as the hydrophobic residues (Figures 1 and 6). This composition, despite the novel arrangement, makes this peptide analogous to others (16). It is well known that the amino acid composition can influence the mechanism of action (7) and perhaps this is the reason why EcDBS1R6 behaved similarly to the positive control melittin, in the experiments probing the mechanism of action (Figures 2 and 3), causing membrane damage (Figure 4).

Originally, EcDBS1R6 was derived from the N-terminus of a mercury transporter protein, which has a signal peptide topology (Figure 1). The signal peptides have similar topologies to transmembrane proteins (13), and therefore, they are able to interact with biological membranes without killing microorganisms (8, 20, 21). The amino acid changes promoted by joker essentially increased the positive net charge of the peptide, which in turn, may have facilitated membrane interaction (Figure 6C). However, increasing the net charge is insufficient to enable the antimicrobial activity, since two other variants of the same parent sequence, EcDBS1R1 and 7, were inactive despite similar charge increase (8). The position of each amino acid in the sequence appears to be a determinant factor for activity (17, 22).

Interestingly, some residues seem to be unnecessary to the activity. The sliding window experiment indicated that the Pro^17^-Trp^19^ residues did not contribute to the activity, since the three C-terminal 10 residue fragments, were not active against *P. aeruginosa* (Figure 5). The core of peptide activity was the N-terminal portion and molecular dynamics indicated that the α-helix was maintained in this region (Figure 6). Indeed, in the classical models of membrane disruption by AMPs, an α-helix formation is often a key element, where for each amino acid that adopts such a conformation there is a free energy change of −0.14 kcal.mol^−1^ (23). Thus the contribution of Ala^12^-Trp^19^ residues is reduced, because they, more frequently, adopt an extended conformation (Figure 6A), which is similar to the NMR structure of EcDBS1R5 variant (12). Besides, considering the original sequence, the signal peptide topology covers the same active region, while the extended region belongs to the mature protein (Figure 1), which indicated that in fact the modifications transformed the signal peptide into a selective agent that might have acted in a detergent-like manner.

## 4 Conclusion

In Nature, the occurrence of gene duplication events and insertions/deletions in addition to cumulative point mutations allows the functional changes in proteins and peptides (24). The Joker algorithm, is capable to enable the antimicrobial activity through a series of mutations and here these mutations were able to transform a signal peptide into an antimicrobial peptide. The use of an algorithm enabled the creation of sequences to fit our needs (7, 10). Currently, there is a great need for new antimicrobial agents, and the peptide EcDBS1R6 was able to kill bacteria of the WHO “priority pathogens” list, indicating a biotechnological potential to develop novel therapeutic agents. In addition identifying the active core allows for the development of small molecules or even the functionalization of non-functional residues. Despite being at an early stage, EcDBS1R6 represents a promising candidate for drug development, mainly for controlling the critical bacteria from WHO list and in future, this peptide could help to solve the bacterial resistance crisis.

## 5 Experimental procedures

### 5.1 Peptide synthesis and mass spectrometry

EcDBS1R6 was synthesized via Fmoc chemistry using a Liberty Blue™ automated microwave peptide synthesizer (CEM Corporation), a PAL-NovaSyn TG resin (Merck Millipore), and a systematic double-coupling protocol. Fmoc-protected amino acids were purchased from Iris Biotech GMBH, and solvents from Carlo Erba. Cleavage and deprotection of the peptidyl resin were achieved by incubation of the resin with an acidic cocktail (95% TFA, 2.5% triisopropylsilane, 2.5% water) for 3 h at room temperature. After removal of the resin by filtration, the crude peptide was precipitated with cold diethyl ether (3000 × g, 15 min, 4 °C), washed with the same solvent, and dried under a stream of nitrogen. The crude peptide was subjected to purification by reversed-phase HPLC on a semi-preparative Nucleosil C-18 column (5 μm, 250 × 10 mm, Interchim SA) using a 20-60% linear gradient of acetonitrile. The homogeneity and identity of synthetic peptides were assessed by MALDI-TOF MS/MS analysis on an UltraFlex III (Bruker Daltonics). The peptide monoisotopic mass was obtained in reflector mode with external calibration, using the Peptide Calibration Standard for Mass Spectrometry calibration mixture (up to 4000 Da mass range, Bruker Daltonics).

### 5.2 Determination of minimum inhibitory concentration (MIC)

The microorganisms used to determine the MIC of EcDBS1R6 included the Gram-positive bacteria *Enterococcus faecalis* (ATCC 29212), *Staphylococcus aureus* (ATCC 25923), and a clinical isolate of methicillin-resistant *S. aureus* (MRSA 7133623); Gram-negative bacteria included *Escherichia coli* (ATCC 25922), *Pseudomonas aeruginosa* (ATCC 27853), *Acinetobacter baumannii* (ATCC 19606), *Klebsiella pneumoniae* (ATCC 13883), a clinical isolate of carbapenem-resistant *K. pneumoniae* (3259271), *Streptococcus pyogenes* (ATCC 19615) and *Listeria ivanovii* (Li 4pVS2). The last two se strains Gram-positive species were cultured in Brain Heart Infusion (BHI) broth, while the others were cultured in Lysogeny Broth (LB). The fungi *Candida albicans* ATCC 90028 and *C. parapsilosis* ATCC 22019 were cultured in Yeast Peptone Dextrose (YPD) medium. MIC was determined in 96-well microtitration plates by growing the microorganisms in the presence of two-fold serial dilution of the peptide, as previously described (25). Briefly, logarithmic phase cultures of bacteria were centrifuged and suspended in MH (Mueller-Hinton) broth to an optical density at 630 nm (OD_630_) of 0.01 (assuming 10^6^ cfu.ml^−1^ for each strain), except for *S. pyogenes*, *L. ivanovii*, *E. faecalis* and *Candida species* which were suspended in their respective growth medium. Microtiter plate wells received aliquots of 50 μl each of the diluted peptide prepared in sterile milliQ water (200 to 1 μM, final concentrations) followed by the addition of 50 μl of the culture suspension. After overnight incubation (18-20 h) at 37 °C (30 °C for fungi), antimicrobial susceptibility was monitored by measuring the change in OD_630_ value using a microplate reader. MIC was determined as the lowest peptide concentration that completely inhibited the growth of the microorganism and corresponds to the average value obtained from three independent experiments, each performed in triplicate with positive (0.7% formaldehyde) and negative (without peptide) inhibition control (26).

### 5.3 Cytotoxic activity

The cytotoxicity of EcDBS1R6 was determined against the human embryonic kidney cell line HEK-293. HEK-293 cells were cultured in DMEM medium supplemented with 10% FBS, GlutaMAX™ 1/100 (v/v), 100 UI/ml penicillin, 100 *μ*g/ml streptomycin, and incubated at 37 °C in a humidified atmosphere of 5% CO_2_. The cell viability was quantified after peptide incubation using a methylthiazolyldiphenyl-tetrazolium bromide (MTT)-based assay (27). Briefly, cells (5 × 10^5^ cells.ml^−1^) were seeded on 96-well culture plates (100 μL / well) and incubated overnight at 37 °C. Cells were then washed with DMEM medium and incubated with 100 μl of EcDBS1R6 (12.5 to 200 μM, final concentrations) during 24 h at 37 °C. Then, 10 μl of MTT (5 mg·ml^−1^) was added to each well and incubated for 4 h in the dark. In living cells, mitochondrial reductases convert the MTT tetrazolium to formazan, which precipitates. Formazan crystals were dissolved using a solubilization solution (40% dimethylformamide in 2% glacial acetic acid, 16% sodium dodecyl sulfate, pH 4.7), followed by 1 h incubation at 37 °C under shaking (150 rpm). Finally, the absorbance of the resuspended formazan crystals was measured at 570 nm. The inhibitory concentration 50 (IC_50_), which corresponds to the peptide concentration resulting in 50% cell death, was determined with GraphPad Prism 5.0 software. Results were expressed as the mean of three independent experiments performed in triplicate.

### 5.4 In vitro *selectivity index calculation*

According to Porto et al. (7), the *in vitro* selectivity index of EcDBS1R6 was calculated using the following equation:

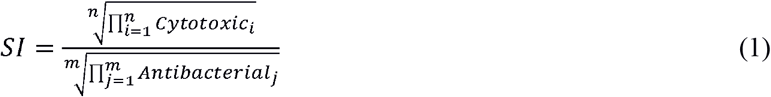

where n is the number of cytotoxicity assays and m is the number of antimicrobial assays with different bacteria. For values higher than the maximum concentration tested, 2-fold the maximum tested value was assumed (e.g. if the value was > 100, it was considered as 200) (28).

### 5.5 Time-kill studies

As previously described (29), exponentially growing bacteria in LB (*S. aureus* ATCC 25923 and *E. coli* ATCC 25922) were harvested by centrifugation, washed and suspended in PBS (10^6^ cfu.ml^−1^, final cell density). After incubation of 100 μl of this bacterial suspension with concentrations of EcDBS1R6 2-fold and 4-fold above the MIC (for *S. aureus* and *E. coli*, respectively), aliquots of 10 μl were withdrawn at different times, serially diluted and spread onto LB agar plates. After overnight incubation at 37 °C, cfu were counted. Two experiments were carried out in triplicate. Negative controls were run without peptide.

### 5.6 SYTOX Green uptake assay

The EcDBS1R6-induced permeation of the cytoplasmic membrane of *S. aureus* ATCC 25923 and *E. coli* ATCC 25922 was measured using the SYTOX Green (SG) uptake assay (30). As previously described (31), exponentially growing bacteria (6 × 10^5^ cfu.ml^−1^) were suspended in PBS after centrifugation (1000 × g, 10 min, 4 °C) and three washing steps. 792 μl of the bacterial suspension was preincubated with 8 μl of 100 μM SG for 30 min at 37 °C in the dark. After peptide addition (200 μl, final concentration 6.25 μM), the fluorescence was measured for 1 h at 37 °C, with excitation and emission wavelengths of 485 and 520 nm, respectively. Three independent experiments were performed, and the results presented correspond to a representative experiment, including negative (PBS) and positive (melittin) controls.

### 5.7 Membrane depolarization

The cytoplasmic membrane depolarization activity of EcDBS1R6 was measured using *S. aureus* ATCC 25923 and *E. coli* ATCC 25922 and the membrane potential sensitive probe DiSC3(5) (3,3’-dipropylthiadicarbocyanine iodide) (32). As previously described (33), exponentially growing bacteria were centrifuged, washed with PBS and resuspended to an OD_630_ of 0.1 (1 × 10^7^ cells.ml^−1^) in the same buffer. 700 μl of bacteria were preincubated with 1 μM DiSC3(5) in the dark for 10 min at 37 °C, and then 100 μl of 1 mM KCl were added in order to equilibrate the cytoplasmic and external K^+^ concentrations. After addition of peptide (6.25 μM), the fluorescence was monitored at 37 °C for 20 min with excitation and emission wavelengths of 622 and 670 nm, respectively. Three independent experiments were performed and the results presented correspond to a representative experiment, including negative (PBS) and positive (melittin) controls.

### 5.8 FEG-SEM imaging

Scanning Electron Microscopy with Field-Emission Gun (FEG-SEM) was used to obtain high-resolution images of *L. ivanovii* (Li 4pVS2) and *P. aeruginosa* (ATCC 27853) treated with EcDBS1R6. Bacteria in mid-logarithmic phase were collected by centrifugation, washed twice with PBS, and resuspended in the same buffer (2 × 10^7^ cfu.ml^−1^). 200 μl of the bacterial suspension were incubated with EcDBS1R6 (12.5 μM) for 1 h at 37 °C. Negative controls were run in buffer without peptide. Cells were fixed with 2.5% glutaraldehyde. FEG-SEM images were recorded with a Hitachi SU-70 Field-Emission Gun Scanning Electron Microscope. The samples (20 μl of inoculum deposited and dried under dry nitrogen on gold plates) were fixed on an alumina SEM support with a carbon adhesive tape and were observed without metal coating. Secondary electron detector (SE-Lower) was used to characterize the samples. The accelerating voltage was 1 kV and the working distance was around 15 mm. At least five to ten different locations were analyzed on each surface, leading to a minimum of 100 single cells observed.

### 5.9 Active core identification

The EcDBS1R6 sequence was submitted to a sliding window system of ten residues, generating ten fragments. These fragments were synthesized in a peptide array designed and synthesized by Kinexus Bioinformatics Corporation (Vancouver, BC). Peptides were produced in a standard mass of 80□μg by using cellulose support in SPOT technology (34). The crude synthetic peptides were obtained from cellulose membrane discs that had already been treated with ammonia gas to release the peptides from the membrane. Peptides were then dissolved overnight in distilled water and subsequently evaluated for the antimicrobial activity.

The antimicrobial activity of the synthesized peptides was evaluated against an engineered luminescent *P. aeruginosa* H1001 (fliC∷luxCDABE) strain in 96-well microplates (35). Aqueous solutions of peptides released from the cellulose spots were diluted two fold in BM2 medium □62□mM potassium phosphate buffer pH 7; 2□mM MgSO4; 10□μM FeSO4; 0.4% (wt/vol) glucose] down the eight wells of a 96-well plate, achieving a final volume of 25□μl in each well. Subsequently, 50□μl of overnight culture of *P. aeruginosa* H1001 (fliC∷luxCDABE) were subcultured in 5□ml of fresh LB media and grown until they reached an OD_600_ of 0.4. This growing bacteria culture was then diluted 4:100 (v/v) into fresh BM2 medium and 25□μl of this diluted bacterial culture were transferred to the microplate wells containing 25□μl of peptide solution. The final peptide concentrations tested ranged from 200 to 3□μg.ml^−1^. The plates were incubated for 4□h at 37□°C with constant shaking at 50□rpm. Luminescence was measured on a Tecan SPECTRAFluor Plus Microplate Reader (Tecan US, Morrisville, NC). The antimicrobial activity was evaluated by the ability of the peptides to reduce the luminescence of *P. aeruginosa*-lux strain compared to untreated cells. The carbapenem meropenem was used as positive control and distilled water was used as a negative control.

### 5.10 Ab initio*three-dimensional structure prediction*

The three-dimensional structure prediction was performed according to Porto *et al*. (8). QUARK *ab initio* modeling server (36) was used for generating the three-dimensional models of the designed sequences. This server was one of the best ranked servers for free modeling in the last four CASP editions and has been successfully used for predicting the structure-function relationship of short peptides, as described earlier (37–39). Because the minimum input length is 20 amino acid residues, glycine residues were added to the C-terminus of sequences of smaller size. After modeling, the additional glycine residues were removed and the structures were submitted to an energy minimization procedure using the GROMOS 96 force field implementation of Swiss PDB Viewer (40), and 2000 steps in the algorithm steepest descent. The models were evaluated through RAMPAGE (41). RAMPAGE checks the stereochemical quality of a protein structure, through the Ramachandran plot, where reliable models are expected to have > 90% of amino acid residues in most favored and additional allowed regions.

### 5.11 Molecular Dynamics Simulations

The molecular dynamics simulations were performed in GROMOS96 43A1 force field using the GROMACS 4 package (42). Structures were simulated in a water/2,2,2-trifluoroethanol (TFE) mixture (50%, v/v). To achieve such a concentration, a ratio of 1:4 TFE to water molecules was used. The environment was constructed using the Single Point Charge water model (43) and a TFE model generated by PRODRG server (44) with reduced charges. The simulations were done in cubic boxes with 0.8 nm minimum distances between the peptides and the limits of the box. Chlorine or sodium ions were added to neutralize the systems with non-zero charge. The water molecule geometry was controlled by the SETTLE algorithm (45). All atomic bonds were constrained using the LINCS algorithm (46). Electrostatic corrections were made by Particle Mesh Ewald algorithm (47) with a 1.4 nm threshold to minimize computational time. The same cutoff was used for van der Waals interactions. The list of neighbors of each atom was updated every 5000 simulation steps of 2 fs. The system underwent an energy minimization using 50,000 steps of the steepest descent algorithm. System temperature was normalized to 300 °K for 100 ps, using the velocity-rescaling thermostat (NVT ensemble), and the system pressure was normalized to 1 bar for 100 ps, using the Parrinello-Rahman barostat (NPT ensemble). Simulations of the complete system were done for 100 ns using the leap-frog algorithm as the integrator. The simulation was repeated 50 times. Molecular dynamics simulation trajectories were analyzed by means of the backbone root mean square deviation (RMSD).

### 5.12 Electrostatic Surface Calculation

The solvation potential energy was measured for the most common structure representative structure at 100 ns of simulation. The conversion of PDB files into “.pqr” files was performed by the utility PDB2PQR using the AMBER force field (48). The grid dimensions for Adaptive Poisson-Boltzmann Solver (APBS) calculation were also determined by PDB2PQR. Solvation potential energy was calculated by APBS (49). Surface visualization was performed using the APBS plugin for PyMOL.

## 6 Acknowledgements

HEK-293 cells were kindly provided by Dr. O. Jean-Jean (UMR 8256, IBPS, FR 3631 SU-CNRS, Sorbonne Université, Paris, France). We thank Dr. S. André (BIOSIPE, IBPS, FR 3631 SU-CNRS, Sorbonne Université, Paris, France) for her assistance in the lab experiments, Dr. C. Piesse (Peptide Synthesis Platform, IBPS, FR 3631 SU-CNRS, Sorbonne Université, Paris, France) for help in synthesis and purification of peptides, and the Mass Spectrometry and Proteomics Platform (IBPS, FR 3631 SU-CNRS, Sorbonne Université, Paris, France) for MALDI-TOF analysis, and also Sandra Casale (Laboratoire de Réactivité de Surface, UMR 7197 CNRS-SU, Sorbonne Université, Paris, France) for her technical assistance in FEG-SEM imaging. The authors acknowledge UPMC/Sorbonne Université for financial support, IMPC (Institut des Matériaux de Paris Centre, FR 2482) and the C’Nano projects of the Région Ile-de-France for SEM-FEG funding. Conselho Nacional de Desenvolvimento Científico e Tecnológico – CNPq for financial support, Coordenação de Aperfeiçoamento de Pessoal do Nível Superior – CAPES, for scholarship, Fundação de Amparo à Pesquisa do Distrito Fedearal (FAPDF) and Fundação de Apoio ao Desenvolvimento do Ensino, Ciência e Tecnologia do Estado de Mato Grosso do Sul (FUNDECT). REWH received funding from the Canadian Institutes for Health Research grant FDN-154287 and holds a Canada Research Chair in Health and Genomics and a UBC Killam Professorship.

## Author contributions

WFP, AL and OLF designed the experiments; LNI, VH and SMR performed and analyzed the *in vitro* experiments; WFP performed and analyzed the computational simulations; WFP wrote the revised; AL, EFH, REWH and OLF revised the manuscript.

